# Phylogenetic Analysis of the Novel Coronavirus Reveals Important Variants in Indian Strains

**DOI:** 10.1101/2020.04.14.041301

**Authors:** Aditi Joshi, Sushmita Paul

## Abstract

Recently classified as a pandemic by WHO, novel Corononavirus 2019 has affected almost every corner of the globe causing human deaths in a range of hundred thousands. The virus having its roots in Wuhan (China) has been spread over the world by its own property to change itself accordingly. These changes correspond to its transmission and pathogenicity due to which the concept of social distancing appeared into the picture. In this paper, a few findings from the whole genome sequence analysis of viral genome sequences submitted from India are presented. The data used for analysis comprises 440 collective genome sequences of virus submitted in GenBank, GISAID, and SRA projects, from around the world as well as 28 viral sequences from India. Multiple sequence alignment of all genome sequences was performed and analysed. A novel non-synonymous mutation 4809C>T (S1515F) in NSP3 gene of SARS-CoV2 Indian strains is reported along with other frequent and important changes from around the world: 3037C>T, 14408C>T, and 23403A>G. The novel change was observed in samples collected in the month of March, whereas was found to be absent in samples collected in January with the respective persons’ travel history to China. Phylogenetic analysis clustered the sequences with this change as one separate clade. Mutation was predicted as stabilising change by insilco tool DynaMut. A second patient in the world to our knowledge with multiple (Wuhan and USA) strain contraction was observed in this study. The infected person is among the two early infected patients with travel history to China. Strains sequenced in Iran stood out to have different variants, as most of the reported frequent variants were not observed. The objective of this paper is to highlight the similarities and changes observed in the submitted Indian viral strains. This helps to keep track on the activity, that how virus is changing into a new subtype. Major strains observed were European with the novel change in India and other being emergent clade of Iran. Its important to observe the changes in NSP3 gene, as this gene has been reported with extensive positive selection as well as potential drug target. Extensive Positive Selection Drives the Evolution of Nonstructural Proteins. With the limited number of sequences this was the only frequent novel non-synonymous change observed from Indian strains, thereby making this change vulnerable for investigation in future. This paper has a special focus on tracking of Indian viral sequences submitted in public domain.

## Introduction

The pandemic of COVID-19 is taking a troll in health and finance sector of all most every country. The Severe Acute Respiratory Syndrome Coronavirus 2 (SARS-CoV2) originated from Wuhan a city in China is mutating while getting transmitted [3]. Development of a potential drug or vaccine is a major challenge. The virus has RNA as a genetic material and it gets mutated frequently. It is very important to understand the evolutionary feature in the virus. Understanding the nature of mutation may help in developing potential drug. Also, it may give direction to understand the next cycle of mutation in the viral genome. Therefore, it is very important to understand the nature of variation in different strains of the novel virus.

In this regard, few studies are conducted to understand the nature of variation in this novel virus. Carmine and Federico [4] made an attempt to understand the heterogeneity in 54 strains of this viral genome. Rahila et al. [16] studied impact of the mutations in the spike glycoprotein function and stability in the novel virus. Changchuan [20] presented a method for effectively genotyping SARS-CoV-2 viruses using complete genomes.

In this report, an analysis of 468 novel and early virus genome is presented. The 440 sequences are from different parts of the world including countries from Europe, USA, Middle East, Asia and 28 sequences submitted from India with 11 sequenced in Iran. The data was downloaded from GenBank, GISAID, and SRA. Data was collected for this study last as of 9th April 2020, with 440 genome sequence of the novel and early strains from the world along with 28 sequences from India. The MEGAX tool [11] has been used to perform the analysis. It is based on Tamura-Nei model and Neighbour-Joining method along with 100 bootstrap. This study adds up a novel non-synonymous mutation 4809C>T (S1515F) in NSP3 gene only observed in strains obtained from India. Various studies revealed that the NSP3 gene harbours many mutations as this gene has been reported with extensive positive selection towards evolution in Betacoronaviruses [18], [7]. Common mutations among different strains were also identified and reported. The sequence containing the mutation was used to build protein model using SWISS-Model. Followed by the usage of the DynaMut tool to study effect of mutation on protein stability. The results indicate the clade of variations in the Indian SARS-CoV2. Also, potential region of mutations in near future along with the effect of non-synonymous mutation at the protein level. This paper represents the initial observations on submitted 28 sequences of Indian viral strains.

## Materials and Methods

Sequence collection: total 468 sequenced strains were selected, 440 from different parts of the world including countries from Europe, USA, Middle East, Asia and 28 sequences submitted from India with 11 sequenced in Iran. Sequences were downloaded from GenBank, GISAID, and SRA project (PRJNA607948). Information of samples used in this study are provided as supplementary material.

SRA project sequences were aligned to human genome assembly (hg19) using HiSat2 [10], to remove the human contamination in viral sequences. Next the unaligned region to hg19 was aligned to SARS-CoV2 ref sequence (NC_045512) using SamTools [13] and HiSat2. Consensus fasta sequence of four viral strains of project were obtained using BcfTools [13].

Multiple Sequence alignment was performed on all 468 sequences using MAFFT (Multiple Alignment using Fast Fourier Transform). Alignment was visualised in JalView 2.11.0. The reported and novel mutation site were extracted to calculate the position specific frequency in all the strains. SWISS-Model was used to build the protein model of fragment containing the mutation. DynaMut was used to predict the 50 possible effect of mutation on protein stability. The phylogenetic tree was constructed 51 with Tamura-Nei model and Neighbour-Joining method along with 100 bootstrap steps 52 using the MEGAX.

## Results

This section describes about the different types of mutations discovered in the analysis along with an analysis of the mutation at protein level.

### The Common Mutations

Genome sequences of 440 viral strains submitted from different parts of the world were collected from GenBank, GISAID and SRA project. In order to investigate the changes in 28 viral strains submitted from India a sequence level and phylogenetic analysis was performed. Multiple sequence alignment of these collected sequences was visualised to collect the positions in Indian strains with mutations.

There has been a report on SNP mutations in SARS-CoV2 genome which majorly demonstrated mutations frequent in European strains [20]. These mutations were analysed in 28 submitted sequences from India. A previous report from India observed unique mutation in spike protein (A930V (24351C>T)), this mutation was identified in one (Sample_2) of the early infected patients’ viral sequence [16]. The most common SNP’s that were observed in this analysis includes the previously reported mutation in leader sequence (241C>T), Coronaviruses includes subgenomic identical 5′ Leader sequence which plays role in viral replication process. This mutation was seen to co-evolved with other three important mutations in gene NSP3: 3037C>T (synonymous), RNA primase: 14408C>T and Spike protein: 23403A>G. In context to Indian strains the early collected samples with travel history to China (Sample_1 and Sample_2) exhibited none of the co-evolved variation (3037C>T, 14408C>T and 23403A>G) with leader sequence variant (241C>T). With recently submitted 11 strain sequences of Indian resident sequenced in Iran (Sample_18 to Sample_28), all the three variants along with leader sequence variant was not observed. In remaining 14 strains (Sample_3 to Sample_16), these four changes were observed as these included (Sample_5 to Sample_9) Italian tourist, one (Sample_3) with travel history to Italy, one (Sample_4) was Indian contact to Italian tourist and 7 (Sample_10 to Sample_16) strains from VERO CCL81 isolate P1. These three along with 241C>T, co-mutations were suggested to increase the transmission [20].

### Mutations in the Different Non-Structural Protein (NSP) Genes

Two of the mutations 3037C>T (F106F) and 14408C>T (P323L) are associated with RNA replication mechanism. The variation reported in NSP3 (3037C>T) lies in N-terminal of the gene. In SARS-CoV gene NSP3 has a critical interaction with Nucleocapsid protein via N-terminal ubiquitin like domain (Ubl), there results suggested that the interaction serves to tether the genome to the newly translated replicase-transcriptase complex at a very early stage of infection [8]. Spike protein has been a hotspot in CoV2 genome since the beginning due to the presence of receptor-binding domain. Contact point to the host includes the domains from S1 subunit of this S-protein [19]. The mutation observed 23403A>G (D614G) lies in tail region of this S1 subunit and before the cleavage site of another subunit called S2 of S-protein.

Talking in terms of RNA replication of SARS-CoV ORF1a and ORF1b (replicase/transcriptase) which includes NSP8 and NSP4 genes have been highlighted. Two linked SNP’s one in NSP4, 8782C>T (S75S) included in codon for Serine residue and other in NSP8, 28144T>C (L84S) included in codon for Leucine residue, according to the population genetic study on 103 SARS-CoV2 genomes has been named as S-type(“CT-bases at two positions”) and L-type(“TC-bases at two positions”) of strains [17]. Strikingly sequence from one (Sample_2) of the early infected persons’ viral genome contains both linked SNP while the other one has absence of both. From other sequences only two (Sample_8 and Sample_9) strains of Italian tourist were presented with 8782C>T change. The other abundant variation observed in Europeans was 26144G>T (G251V) in NSP3 gene and three consecutive mutations observed the most in US strains 28881G>A (R203K), 28881G>A (R202R), 28882G>A (G204R). Surprisingly, this 26144G>T change was not observed in any of the strains, including the strains from Italian tourists. Three consecutive variations were observed in only one (Sample_2) of the early strain that also contained both S-type and L-type linked SNP.

Recent strains (Sample_18 to Sample_28) submitted from Indian resident sequenced at Iran had mutations in other positions then mentioned above. Position 884C>T (Figure 1) change in NSP2 gene was observed in 9 strains out of 11 (Sample_18 to Sample_25, Sample_27 and Sample_28), these variations were observed in 2 strains (GISAID Id: EPI ISL 416541 and EPI ISL 416458) from Kuwait and 1 strain (GISAID ID:EPI ISL 417444) from Pakistan Gilgit region. Out of two from Kuwait one had travel history to Iran and other was sample from Quarantine center. Also Pakistan strain collected from sample had a travel history to Iran. Second variant 1397G>A (Figure 1) reported in preprint [14] from NSP2 gene was observed in all Indian strains (Sample_18 to Sample_28) from Iran along with same 2 (GISAID Id: EPI ISL 416541 and EPI ISL 416458) strains from kuwait, 1 Pakistan Gilgit strain (GISAID ID:EPI ISL 417444) and additionally in one strain (GISAID IF: EPI ISL 417521) from Taiwan with no information about travel. Variant 8653G>T (Figure 1) in NSP4 gene was observed in 9 strains (Sample_18 to Sample_25, Sample_27 and Sample_28), along with two strains (GISAID Id: EPI ISL 416541 and EPI ISL 416458) from Kuwait. The reported variant 11083G>T (Figure 1) of NSP6 was presented in all strains (Sample_18 to Sample_28) which had a reported frequency of 115 from [20]. Another frequent mutation 28688T>C (Figure 1) in ORF9, was observed in all Indian strains (Sample_18 to Sample_28) sequenced in Iran. This variation was observed in above mentioned two Kuwait strain (GISAID ID:EPI ISL 417444) and one strain from Taiwan (GISAID IF: EPI ISL 417521). The variant 29742G>T (Figure 1), was observed in 3’UTR region. Base change observed at this position G>T; was identified in 2 strains from Kuwait (GISAID Id: EPI ISL 416541 and EPI ISL 416458) and 1 strain from Taiwan (GISAID IF: EPI ISL 417521), whereas two strain sequences from US (MT106052.1 and MT263386.1) presented G>A at this position.

**Figure 1.**
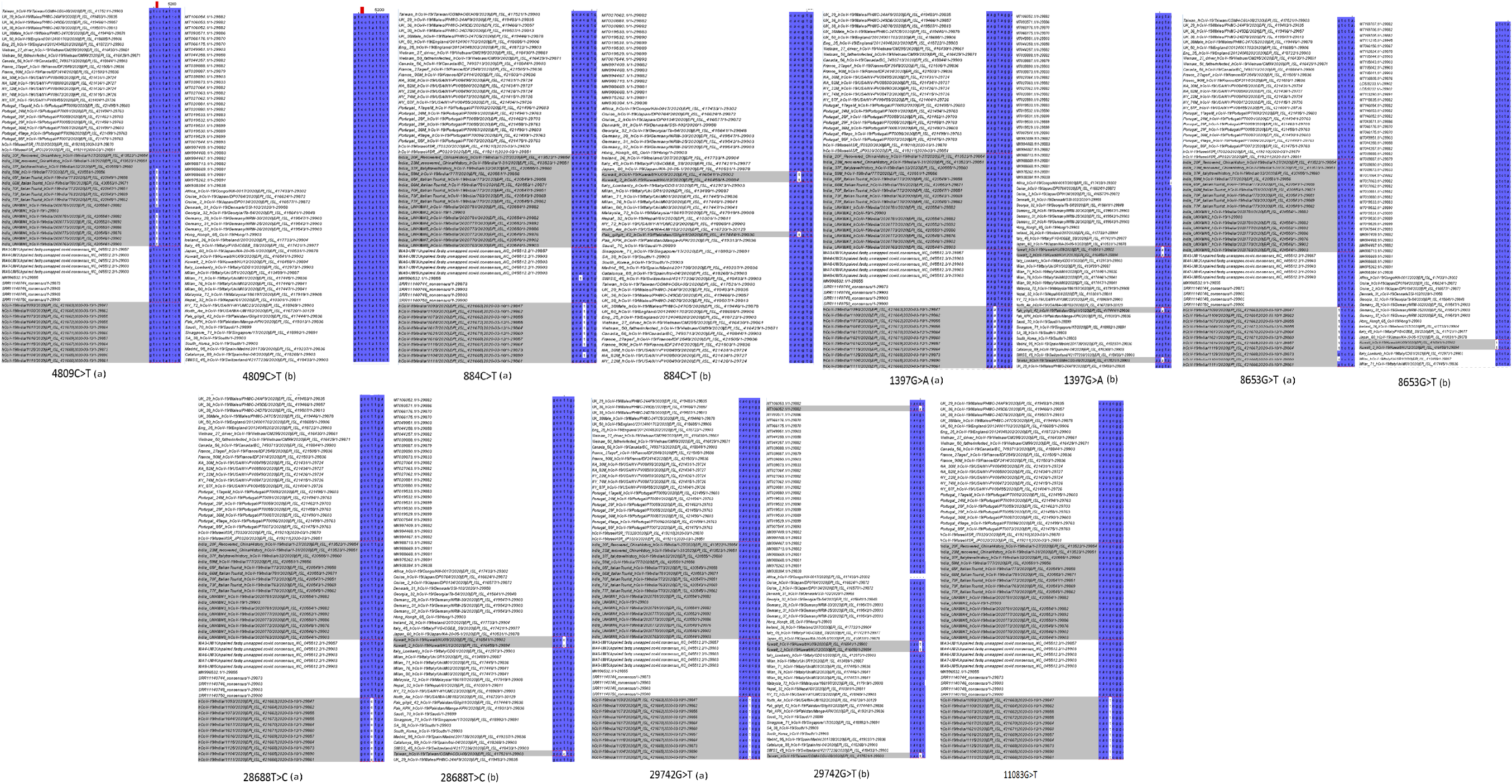
Sequence alignment and variants at different positions

### The Novel Stabilising Mutation in the NSP3 Gene

There was one novel mutation observed in sequences from India in NSP3 gene 4809C>T (S697F) (Figure 1) a non-synonymous change. This observed mutation is close to the reported synonymous variation 3037C>T (F106F). There were 17 strains (Sample_4 to Sample_9 and Sample_11 to Sample_16) which got sequenced in India, out of these 12 strains including sequences from 5 (Sample_5 to Sample_9) Italian tourist, 6 (Sample_11 to Sample_16) Vero CCL81 isolate P1 and one (Sample_4) from sample who was reported to have Indian contact to Italian tourist were observed with this change. Besides the personal point mutations in one or two strains, this novel mutation was observed as frequent among the strains sequenced in India. Interestingly this variant was not found in any European, US or other strains included in this study.

SARS-CoV2 has been reported to have second highest identity (82.30%) with SARS-COV. To identify the possible role and relevance of the region observing the mutation, sequence alignment of NSP3 protein from SARS-CoV and SARS-CoV2 was performed. NSP3 gene along with NSP4 are the key players for membrane rearrangement in form of Double Membrane Vesicles (DMV) upon infection [2]. Report on role of mutations in important glycosylation sites in NSP4 gene in SARS-Cov have been observed and were indicative of localisation problems with other NSP protein like NSP3 [5]. As per this there will be a requirement to observe the site of novel change for its impact on NSP3 gene interactions, which may even serve as a potential candidate for therapeutics, depending upon the mutational effect shown in NSP4 [5]. The other predictive outcome can come as benefit to virus in terms of interaction.

In [8], the authors reported domain organisation of NSP3 gene from SARS-CoV. The mutation was identified in SARS-Unique Domain (SUD). The patch of 162 amino acid near the mutation was observed to align in SARS-CoV’s SUD-C subdomain, a frataxin-like domain. Till now the subdomains in SUD have not been identified in SARS-CoV2. The role of this domain involves binding to single-stranded RNA and recognize purine bases more strongly than pyrimidine bases [9]. Though the very crucial role of SUD-C has not been recognised in SARS-CoV but in few reports SUD-C has been shown to have RNA-binding ability, and concerted actions by the subdomains of SUD have been proposed to ensure specific RNA binding [9].The complete deletion of the SUD-C domain within the context of a SARS-CoV was found to have large reduction of RNA synthesis, but some basal RTC activity remains [12]. PDB blast confirmed the presence of mutation in SUD domain of SARS-CoV with 74% identity to 2KAF (PDB-ID). Using Swiss model this particular domain was modelled for Reference Wuhan sequence (NC 045521)and the change was observed by insilco protein stability tool DynaMut [15]. The change was predicted to be stabilising by DynaMut and destabilising by 3 others. The DynaMut is more elaborative prediction tool. So far a destabilising mutation in NSP3 gene of SARS-CoV2 has been indicative of only differentiation from SARS-CoV [1].

Figure 1 contains MSA alignment of the novel non-synonymous mutation 4809C>T and few mutations from Iran samples.

## Discussion

The novel Coronavirus, being identified as a global threat have been the prime focus of research and investigation around the globe. Since its outbreak in Wuhan (China) it has taken up several changes to keep the survival high. There are reports by several groups on mutations observed in the strains from different countries [16] [20] [16]. Here, in this study, the primary focus is on strains submitted by India. The country is suspected to have more number of positive cases than actually reported and also, there is increase in the number of cases despite the lockdown implemented by the government. With increasing number of cases the virus might have incorporated some new changes to enhance its transmission and virulence. Hence, it becomes important to interrogate the strains submitted to observe the similarities and changes from other strains in the world.

There are some interesting results from this sequence observation analysis. Two of the early collected samples (Sample_1 and Sample_2) with travel history to China were sampled within the gap of 03 days. They were observed with none of the European reported frequent variations. Interestingly one (Sample_2) of them had incorporated the change in both linked SNP’s (8782C>T and 28144T>C) and one could suspect that he might have been infected with the S-type as well as L-type of strain. This is the second report of such event, there has been one more case reported from the US [17]. The cause of this is unclear, but there is a possibility as per previous report that patient might have infected multiple times. This same infected person has another notable change of three consecutive positions (28881G>A, 28882G>A, and 28883G>C) reported mainly in the US. To an extent it clears the doubt that person might have contracted multiple strains (Asian and American), thereby carrying both S and L types of strain along with the US reported mutation. L-type was found to be more aggressive strain before Jan 2020 [17]. People who might have contracted this patient would have been at higher risk of infection. In phylogenetic tree analysis these two strains could be observed without branching from the ancestral strain and in vicinity to reference strain sequence (NC 045512) Figure 2.

**Figure 2.**
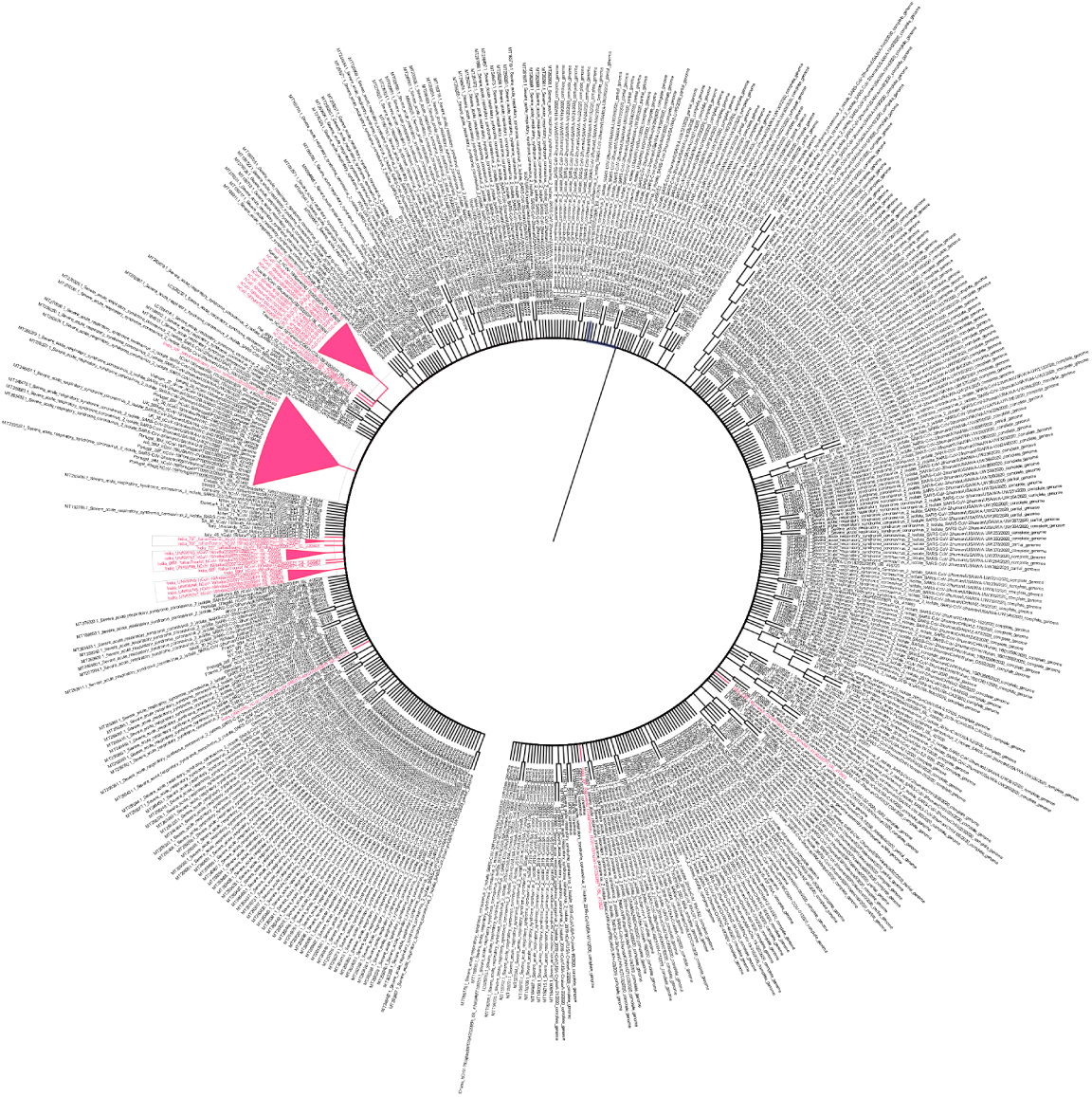
Phylogentic Tree representing the 468 sequences. Indian strains are high-lighted in pink.

Sequences submitted recently from 11 strains (Sample_18 to Sample_28) of Indian citizens sequenced at Iran gave a different picture. These strains does not include any of the reported frequent variant circulating in the world. This clears that these are all Wuhan-like strains with some potential other changes contributing towards the diversity of the virus [6]. This diversity can be associated with fitness or transmission by viral genome. Among the 5 frequent mutations observed in these strains, 2 were reported mutation (11083G>T and 1397G>A, [20]. Divergence from the main strain can be easily pointed out in the constructed phylogenetic tree (Figure 2) with additional root node. These variants could not be observed in sequences submitted from Iran as they had sequenced mainly the nucleocapsid protein.

The novel mutation 4809C>T observed even with this small number of strains sequenced in India with varied travel history and ethnicity. The observed mutation is non-synonymous change observed in 1.7Kb region of NSP3 region, with other being 3037C>T. Indeed the variant is absent in any other strain included in this study along with Europeans. This change has occurred in strains obtained from all the Italian tourists (Sample_5 to Sample_9), the person (Sample_4) who had contracted with Italian tourist and 6 (Sample_11 to Sample_16) of the Vero CCL81 isolate P1 sequenced in India. It seems to be absent in the strain sequenced from patient with travel history to Italy.

One probable explanation to this novel non-synonymous change could be related to its increased transmission rate since it is being observed to be incorporated in combination with already co-existing mutations [20]. The mutation was predicted with stabilising effect, hence no change at secondary structure level was observed Figure 3.

**Figure 3.**
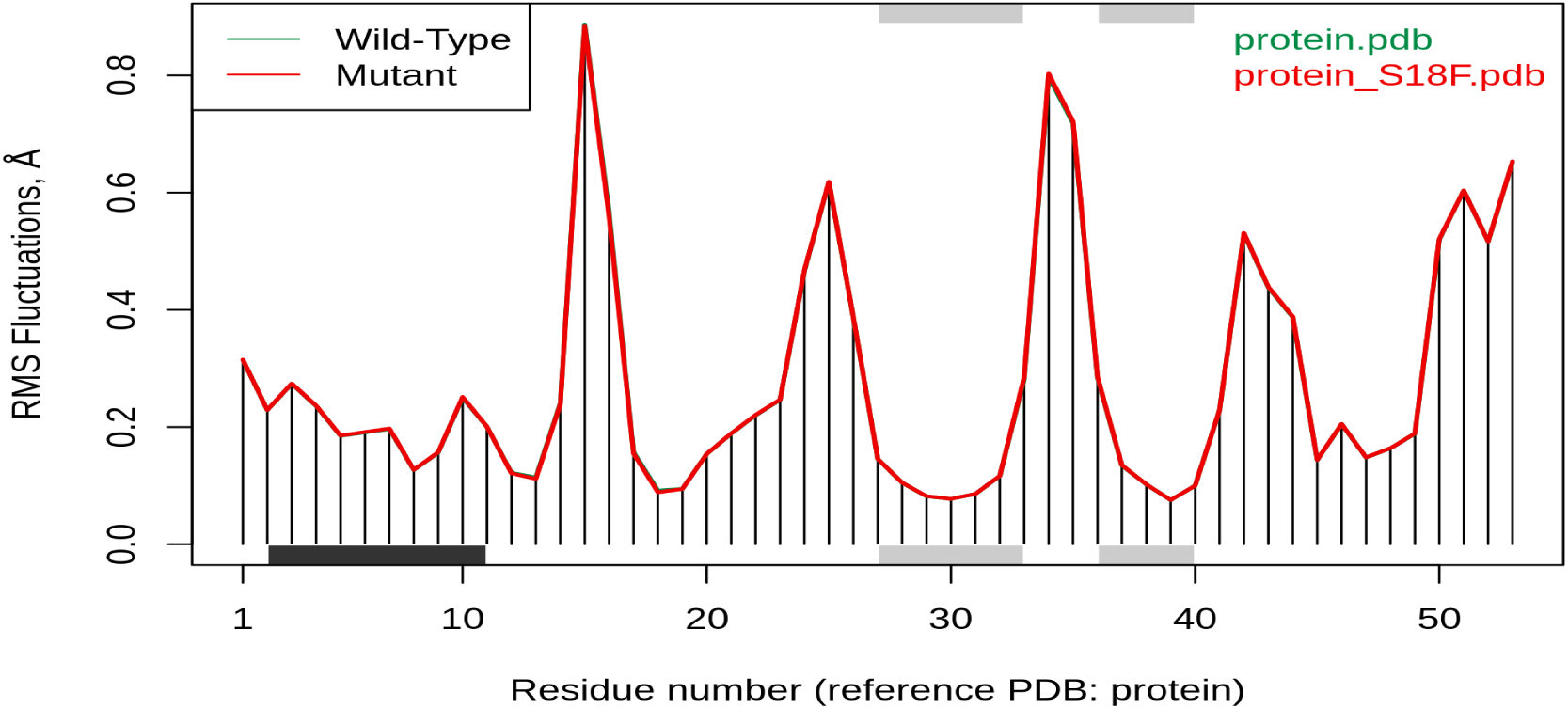
Fluctuation plot predicted by DynaMut comparing the secondary structure in NSP3 submitted domain in wild-type and mutant

Interaction experiments or insilico docking might help to understand its effect more clearly. Adaptive evolutionary changes in NSP3 gene have been discerned with betacoronaviruses, which might contribute towards the host-range or other viral phenotypes [7]. In a recent study [18] on analysing the possible drug targets through computational methods have highligthed the range of chemical and natural products with anti-bacterial and anti-inflammatory properties have high binding affinity to NSP3c (SUD-domain) along with NSP1 and ORF7. Among the few drugs they reported ritonavir, lopinavir and mostly the anti-viral drugs binds with NSP3c. Though they had the limitations and false positives also, but its nonetheless important to consider this domain as potential drug target against SARS-CoV2. This change could be viewed as potential next change a virus might have incorporated to form a new subtype in India as a response to positive selection. Second probable reason could be that the other domains from SUD-SARS-CoV (SUD-N and SUD-M) have been reported to have effect on their binding with G-quadruplexes in human host upon mutations [12]. Thereby change in this SUD-C might have been beneficial, not affecting much the response of virus against host but providing some kind of fitness with the replication or transcription mechanism. As mentioned earlier SUD-C and combination of SUD-MC are involved in RNA binding process [9]. Further biochemical validation or insilico docking analysis are required to observe the effect of this change on RNA-binding.

With earlier reported important mutations 241-C>T, 3037C>T, 14408C>T, 23403A>G this additional novel mutation has been added 241C>T, 3037-C>T, 14408C>T, 23403A>G, 4809C>T presented in Figure 4.

**Figure 4.**
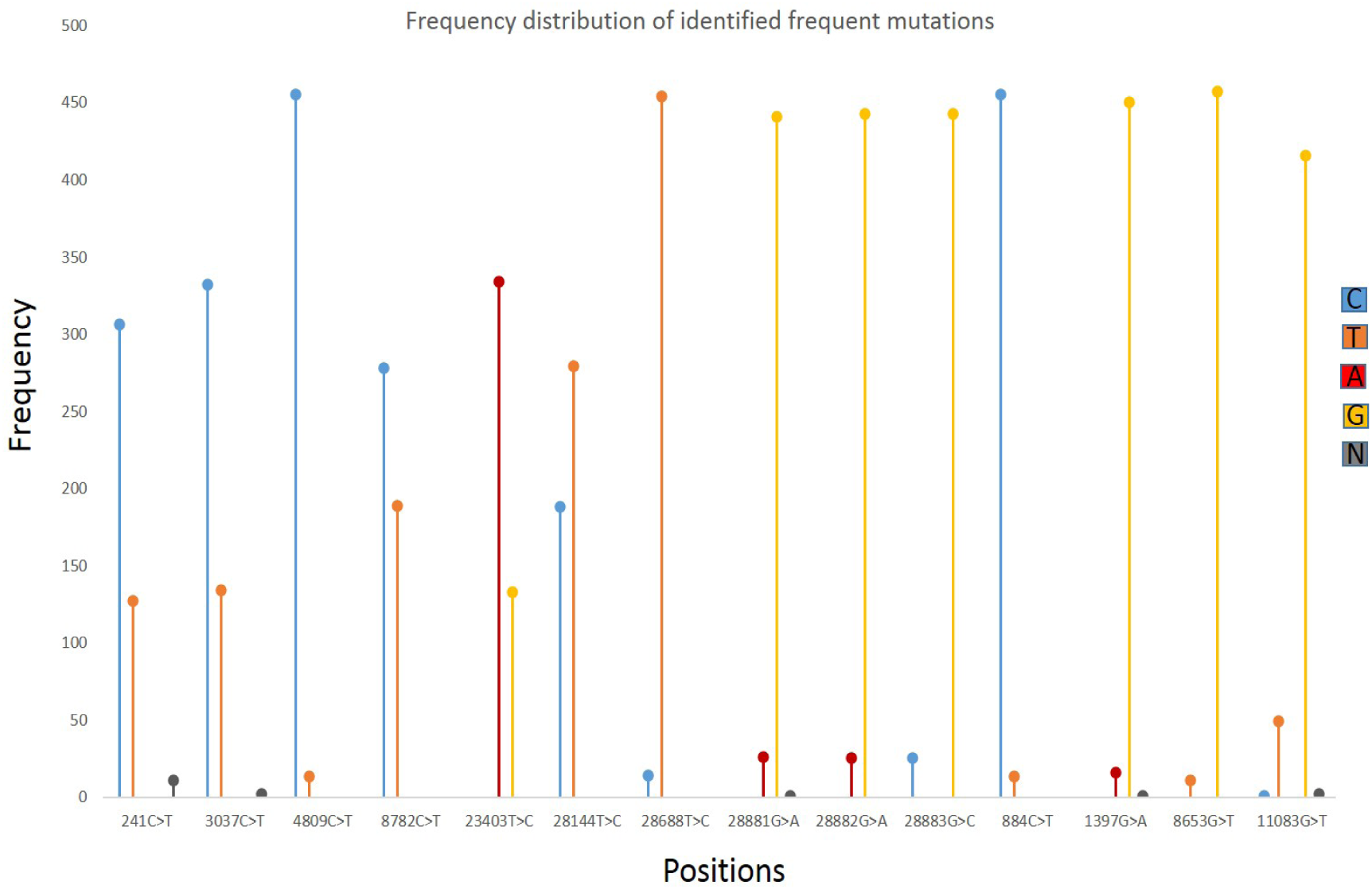
Frequency plot to highlight the changes at frequent positions reported earlier

As discerned with reported frequent mutations in Europeans, the inference could be made that these mutations could be correlated to efficiency of viral transmission, since Europe has emerged with most adverse effects by the virus. The virus have been reported with slow changes and thereby found to be circulating in few major forms. Looking at the data from Indian strains it has majorily the European strain with the novel change observed in this study. The other subtype observed was the Indian citizens sequenced in Iran. In a phylogenetic tree analysis (Figure 2) it was observed that strains containing this novel mutation were branched together and separate clad of citizens came from Iran.

Novel incorporation of one more mutation could be associated with its transmission in Indian population. This change might add up as new member to the group of number of changes the virus has taken in course of evolution in India since the first Italian case identified in Mar 2020. Until now no strain has been declared more lethal in comparison with other, but transmission has been the prime observation, which to an extent could be controlled by social distancing mechanism. This speculation is being made from the very early observations and based on what has been the mutational trend in and around the world. The limitation of this study is the less number of sequences submitted from India. There is an emergent need of more public data sharing from Indian community to track new subtypes of virus, impact of the changes it is incorporating and where the hotspot for therapeutics lies. India is observing the rapid increase in number of cases despite of country lockdown, it makes the analysis of viral strains more crucial. Besides the limitation this study adds up a position to be interrogated in future with more number of sequences and biochemical assays in order to understand its importance in viral machinery.

## Acknowledgments

Authors would like to thank Dr. Mohammed Faruq, Senior Scientist, CSIR-IGIB, Delhi for his valuable support. This work is partially supported by the seed grant program of the Indian Institute of Technology Jodhpur, India (grant no. I/SEED/SPU/20160010).

